# Inhibition of excitatory synaptic transmission alters functional organization and efficiency in cortical neural networks

**DOI:** 10.1101/2023.07.05.547785

**Authors:** Janelle S. Weir, Ola Huse Ramstad, Axel Sandvig, Ioanna Sandvig

## Abstract

Fundamental neural mechanisms such as activity dependent Hebbian and homeostatic neuroplasticity are driven by balanced excitatory – inhibitory synaptic transmission, and work in tandem to coordinate and regulate complex neural network dynamics in both healthy and perturbed conditions. These neuroplasticity processes shape neural network activity, as well as structural and functional aspects of network organization, information transmission and processing. While crucial for all aspects of network function, understanding how the brain utilizes plasticity mechanisms to retain or regain function during and after perturbation is often challenging. This is because these processes occur at varying spatiotemporal scales simultaneously across diverse circuits and brain regions and are thus highly complicated to distinguish from other underlying mechanisms. However, neuroplasticity and self-organizing properties of the brain are largely conserved in *in vitro* biological neural networks, and as such, these networks enable us to investigate both structural and functional plasticity responses to perturbation networks at the micro and mesoscale level. In this study, we selectively silenced excitatory synaptic transmission in *in vitro* neural networks to investigate the impact of the perturbation on structural and functional network organization and resilience. Our results demonstrate that selective inhibition of excitatory transmission leads to transient de-clustering of modular structure, increased path length and degree in perturbed networks. These changes indicate a transient loss of network efficiency; with the network subsequently reorganizing to a state of increased clustering and short path lengths following recovery. These findings highlight the remarkable capacity of neural networks to reconfigure their functional organization following perturbation. The ability to detect and decode such processes as they evolve highlights the robustness of our models to investigate certain dynamic network properties that are often not accessible by *in vivo* methods.

## 1 Introduction

Complex self-organizing systems like *in vitro* neural networks spontaneously acquire structural and functional organization through dynamic activity independent (Goodman and Shatz 1993) and activity dependent (Kater, Davenport et al. 1994, Kirwan, Turner-Bridger et al. 2015) interactions between constituent elements of the network. During network development, structural connectivity is initially characterized by a dense meshwork of neurite connections and synapse overproduction which is subsequently sculpted by experience-driven synaptic pruning and connection refinement (Ivenshitz and Segal 2010, Fauth and Tetzlaff 2016, Millan, Torres et al. 2018). These processes occur concurrently with the maturation of excitatory and inhibitory synapses (Huang 2009), which have pivotal roles in initiating, regulating, and balancing the neural activity within these networks. While excitatory synapses enhance signal transmission, inhibitory synapses act as regulators, effectively controlling and modulating overall network activity levels. The precise coordination between these two types of synapses is crucial for establishing a well-functioning and adaptable neural network (Zhang and Sun 2011). As structural connectivity becomes more refined, these synapses mature and optimize their function, ultimately contributing to the establishment of efficient and effective neural circuitry (Najafi, Elsayed et al. 2020, Sukenik, Vinogradov et al. 2021) manifesting through the emergence of complex network activity such as network bursts and synchrony (Ben-Ari 2001, Opitz, De Lima et al. 2002, Wagenaar, Pine et al. 2006, Chiappalone, Vato et al. 2007).

A key characteristic of self-organizing neural networks is the progressive advancement from a less organized state to the formation of complex topologies and functional hierarchies over time (Karsenti 2008, Prokopenko 2009). The neuroplasticity processes involved in self-organization also dictate network wiring and, by large, information transmission and processing across ordered neural networks (Rubinov, Sporns et al. 2009). Neurons involved in the execution of a specific function tend to cluster together in specialized modules, for example, cells with the same eye preference grouped into ocular dominance columns in the visual system (Hubel and Wiesel 1969), are typically highly interconnected with each other. This organization is ubiquitous across neural networks, where high local clustering of neurons combined with rapid information transmission within and between clusters have been shown to have important implications for functional processing and efficiency (Bassett and Bullmore 2006, Bullmore and Sporns 2009, Meunier, Lambiotte et al. 2010). These modules perform segregated processing facilitated by dense short-range intra-module connections (edges) (Kaiser and Hilgetag 2010, Klinshov, Teramae et al. 2014, Okujeni, Kandler et al. 2017), with global integration with the rest of the network by few long-range inter-module edges (Bullmore and Sporns 2012, Perinelli, Tabarelli et al. 2019). Thus, communication efficiency in the network is inversely proportional to the distance or number of edges (path length) separating processing modules (Kaiser and Hilgetag 2006, Sporns 2013, Sporns 2018).

In complex systems, network resilience to perturbation also plays a critical role in information processing, since failure in one part can trigger cascades of failures throughout the network (Albert, Jeong et al. 2000, Ivanov and Bartsch 2014). In disorders such as post-traumatic stress disorder (PTSD) (Suo, Lei et al. 2015), schizophrenia (Liu, Liang et al. 2008), epilepsy (Li, Chen et al. 2020) and during the prodromal stages of Alzheimer’s disease (AD) (Pereira, Mijalkov et al. 2016), aberrant alterations in path length, clustering and modularity are highly correlated with pathophysiology progression, duration and / or severity. Nevertheless, neural networks can often compensate to maintain function despite any external perturbations or internal fluctuations. Much of this resilience stems from activity-dependent neuroplasticity, which is essential for the brain to function effectively, adapt to changing environments and protect itself against damage (Nakamura, Hillary et al. 2009, Overman and Carmichael 2013). However, since network topology, synaptic transmission and neuroplasticity responses are inextricably linked, quantifying their interdependence in healthy and perturbed conditions in the brain still presents major challenges. Consequently, there are unanswered questions regarding the implications of disrupting synaptic activity on global aspects of network organization and resilience. Specifically, it remains unclear whether the network can maintain its structural and functional organization during perturbation or if it undergoes reorganization, potentially leading to more efficient or less efficient functioning.

For this investigation, we expanded on our previous work using hM4Di designer receptors exclusively activated by designer drugs (DREADDs) (Urban and Roth 2015, Ozawa and Arakawa 2021) to selectively target and inhibit excitatory synaptic transmission in *in vitro* rat cortical networks interfaced with microelectrode arrays (MEAs) (Weir, Christiansen et al. 2023). In our previous work (Weir, Christiansen et al. 2023), we found that selective inhibition significantly influenced the functional electrophysiological dynamics in perturbed networks, which manifested as increased network burst rate, higher fractions of spikes in bursts, and increased synchrony. In the present study, perturbation resulted in transient but significant de-clustering of modules within the networks, with a decrease in clustering coefficient and small-worldness concomitantly with increased path length. These structural changes implied a shift to a random network organization, effectively creating a less optimized and less efficient network.

## 2 Materials and methods

### 2.1 Culture of cortical networks on microelectrode arrays and AAV transduction

Neuronal cultures of primary rat (Sprague Dawley) cortex neurons (Cat. No: A36511) were thawed and co-cultured with 15% primary rat cortical astrocytes (Cat. No: N7745100) both obtained from ThermoFisher Scientific (US). A small drop (80 μl) of the cell suspension containing about 60 × 10^4^ cells were plated on the active area of PDL + Geltrex precoated complementary metal-oxide semiconductor (CMOS)-based high density multielectrode array (HD-MEA) (3Brain GmbH, Switzerland). Cells were incubated in a humidified incubator (5% CO_2_, 37 °C) for 6 hours to allow attachment, then the wells were filled with 1 mL Neurobasal™ Plus Medium supplemented with 2% B-27 Plus supplement and 0.5% GlutaMAX™ all from ThermoFisher Scientific. The day of plating was designated as day 0 and 50% media changes were carried out every 2-3 days. At 7 days in vitro (DIV), the cells were transduced with a vector encoding hM4Di-CaMKlla-DREADDs according to the protocol established in our previous paper (Weir, Christiansen et al. 2023). In short, 80% of the cell media was removed from the cultures and a drop containing 6 × 10^7^ AAV viral particles encoding experimental hM4Di-CamKlla-DREADDs was added to the cells. The cultures were gently agitated for 30 seconds to distribute the viral particles in the media and then incubated for 8 hours. Afterwards, each culture was topped up with fresh Neurobasal Plus cell media and incubated for an additional 40 hours. After the incubation period, 50% media changes were carried out every second day as scheduled. The vector encodes mCherry which is a bright red fluorescent protein tag that makes it possible to visualize results soon after transduction.

### 2.2 Immunocytochemistry

At 14 DIV, samples on Nunc™ Lab -Tek™ chamber slides (Cat. No. 154534PK) were fixed with 4% Paraformaldehyde (PFA) for 20 minutes, then permeabilized with a buffer of 0.03% Triton X-100 and 5% goat serum diluted in DPBS for 2 hours at room temperature. Following blocking, antibodies at the indicated concentrations **(Table 1)** were added in a buffer of 0.01% Triton X-100 and 1% goat serum in DPBS. Nuclei were stained with Hoechst (bisbenzimide H 33342 trihydrochloride, 14533, Sigma-Aldrich, 1: 5000 dilution). Samples were washed, mounted on glass cover slides with anti-fade fluorescence mounting medium (ab104135, Abcam) and imaged. All sample images were acquired using an EVOS M5000 imaging system (Invitrogen, ThermoFisher Scientific). Images were processed using Fiji/ImageJ and Adobe Illustrator 2020 version: 24.0.0.

**Table 1.** Overview of primary and secondary antibodies, species, and concentration

| Primaries |  |  | Secondaries |  |  |
| --- | --- | --- | --- | --- | --- |
| Markers | Catalogue # | Concentration | Fluorescent | Catalogue # | Concentration |
| Ck mCherry | Ab205402 (Abcam) | 1:1000 | Goat-anti-Chicken<br>AlexaFluor 568 | Ab175477<br>(Abcam) | 1:1000 |
| Ms Calmodulin<br>(CaMKII) | MA3-918<br>(Invitrogen) | 1:250 | Goat-anti-Mouse<br>AlexaFluor 568 | A11019<br>(Invitrogen) | 1:1000 |
| Ms NMDAR1 | 32-0500 (Invitrogen) | 1:100 | Goat-anti-Mouse<br>AlexaFluor 647 | A21236<br>(Invitrogen) | 1:1000 |
| Ms GABA | Ab86186 (Abcam) | 1:250 | Goat-anti-Mouse<br>AlexaFluor 488 | A11001<br>(Invitrogen) | 1:1000 |
| Ms TUJ | Ab78078 (Abcam) | 1:500 | Goat-anti-Rabbit<br>AlexaFluor 568 | A11011<br>(Invitrogen) | 1:1000 |
| Rb Glutamate<br>decarboxylase<br>(GAD65/67) | Ab183999 (Abcam) | 1:100 | Goat-anti-Rabbit<br>AlexaFluor 647 | A21244<br>(Invitrogen) | 1:1000 |
| Rb Map2 | Ab32454 (Abcam) | 1:250 | Goat-anti-Rabbit<br>AlexaFluor 488 | A11008<br>(Invitrogen) | 1:1000 |
| Rb Glial fibrillary<br>acidic protein<br>(GFAP) | Ab278054 (Abcam) | 1:500 |  |  |  |

### 2.3 Deschloroclozapine (DCZ) activation of hM4Di DREADDs in neural networks

Networks for DCZ treatment were transiently inhibited once per day at 25, 26 and 27 DIV for 2 hours each time. The novel synthetic ligand deschloroclozapine (DCZ; 10 μM) was used to activate the DREADDs receptors to induce synaptic silencing in excitatory neurons (Bjorkli, Ebbesen et al. 2022, Weir, Christiansen et al. 2023). On the days of treatment, DCZ diluted in cell media was added to 20% media volume in the wells for a final DCZ concentration of 10 μM. Networks were incubated for 2 hours. Thereafter, DCZ was washed out by doing 3 × 80% media changes using fresh media. For each recording, samples were placed in the recording head stage for 10 minutes before starting data acquisition to reduce noise variations due to disturbances caused by moving the cells from the incubator (see **Figure 1** for a workflow). Electrophysiological data was recorded for 15 minutes.

**Figure 1.**
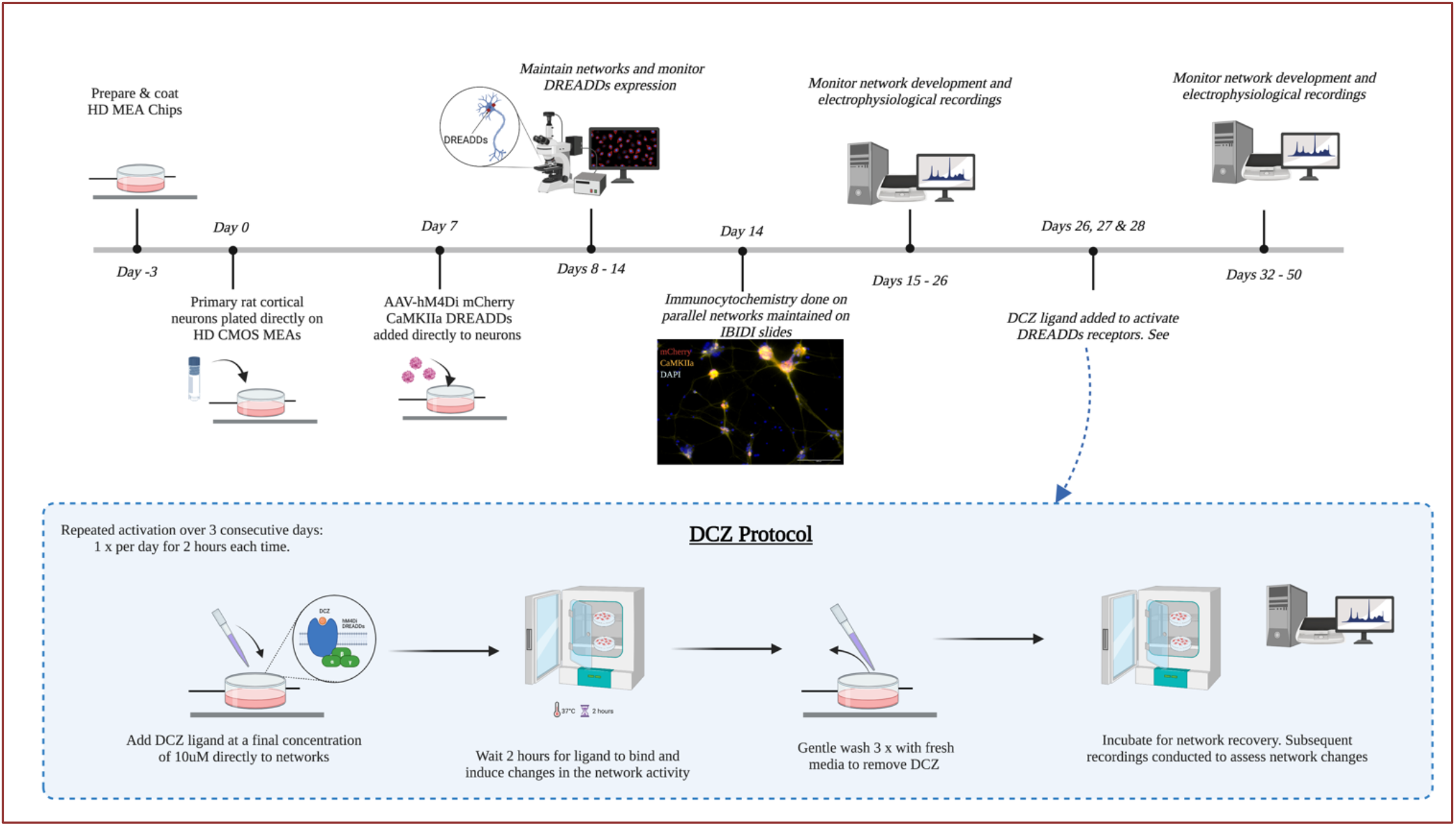
Workflow of experimental procedures. Timeline of experiment from preparation of HD MEAs and throughout the lifetime of the network. The DCZ protocol used for selectively perturbing DREADDs networks is illustrated in the bottom panel. *Figure created in Biorender.com*.

### 2.4 MEA setup and electrophysiological recording

Electrophysiological recordings were performed using the BioCam Duplex 3.0 platform with complementary metal-oxide semiconductor (CMOS) chips. Two models of CMOS chips (Arena and Accura) were used in this study due to availability (3Brain GmbH, Switzerland). The chips are integrated with 4096 square microelectrodes (21 μm x 21 μm) that are arranged in a 64 × 64 square grid on a seeding area of 2.67mm x 2.67mm (Arena) or 3.8mm x 3.8mm (Accura). The electrodes are aligned with 42 μm pitch (Arena) or 60 μm pitch (Accura) between electrodes. Recordings were done at a sampling rate of 20kHz, with a low pass cut off frequency at 100Hz and a high pass cut off frequency of 200 Hz. The raw data was visualized and preprocessed using the BrainWave software v5.1. Preprocessed data was analyzed offline in MATLAB.

### 2.5 Preprocessing: Spike detection and spike sorting using BrainWave 5 (3Brain GmbH, Switzerland)

Spike detection was performed using the Precise Timing Spike Detection (PTSD) algorithm which facilitates reliable and precise identification of spike events. Active channels were defined with a mean firing rate (MFR) of >0.10 spikes/s at a threshold of 7 x standard deviations (std) of the signal’s biological and thermal noise. The peak lifetime period of spikes was set at 2.0 ms with the refractory period set at 1.0 ms. Spike sorting was performed using a K-Means & Silhouette clustering algorithm. Principle component analysis was used for feature extraction with the minimum number of spikes per cluster set at 2, and the maximum at 3. Outlier thresholds were set at 2 and all outliers and duplicate units were discarded. The duplication detection window was set at 0.1 ms with a threshold of 40%.

### 2.6 Network functional connectivity analyses

Connectivity data was analyzed offline using MATLAB (2021b). Functional connectivity was detected using Pearson cross-correlation. Prior to analyzing cross correlation, all potential connections were tested using Spearman’s rank correlation and a bin size of 100 ms. Highly significant (p < 0.001) and strongly correlated (r > 0.01) connections were then further assessed using normalized Pearson cross correlation. For cross correlation, a bin size of 1 ms and a maximum lag of ±100ms was used and significant (p< 0.1) correlations were retained. Correlations of lag 0 were discarded. The weight of a connection in the adjacency matrix was set as the maximum normalized correlation coefficient within the specified lag. A final filtering step was performed to remove connections between distal electrodes with a biologically implausible lag, specifically exceeding 1 mm/ms.

Modularity was determined using functions from the Brain Connectivity toolbox (Rubinov and Sporns 2010) and the Community detection toolbox (https://github.com/mmitalidis/ComDetTB). For each adjacency matrix, the Louvain method of modularity detection (Blondel, Guillaume et al. 2008) was applied 100 times with a gamma of 1 and absolute weights. Each partitioning was then assessed for cluster validity using the node membership criterion as defined in the Community detection toolbox. The partition with the highest cluster validity, that is the strongest internal edge consistency for each module, was then selected. The Modularity Q values indicate the confidence of accurate network subdivision into modules. Modularity can either be positive or negative, with positive values indicating the possible presence of a community structure and negative values or zero indicating an undivided network (Newman 2006). Partitions with modularity Q greater than 0 indicate a community structure and > 0.1 indicate a strong community structure (Newman 2006). In our results, partitions with modularity Q < 0.01 were discarded as too weakly clustered to qualify as a module.

For each module and giant component, the characteristic path length, clustering coefficient and participation coefficient (Guimerà and Nunes Amaral 2005) was detected using the Brain connectivity toolbox. The participation coefficient measures whether a node only interacts with nodes in its own module or if it shares edges with nodes from multiple modules. Path lengths were set as 1 minus the weight to account for stronger correlations being indicative of shorter paths. The intermodular path length describes the average distance between all nodes from one module to all nodes in another module. The clustering coefficient was calculated using binarized edges and describes the property of a node in the network and how well connected the neighborhood is. Mean degree describes the average number of connections that a node has to other nodes in the network. Small-world propensity (SWP) quantifies the extent to which a network displays small-world characteristics while accounting for variations in network density, and was calculated using the methods described in (Muldoon, Bridgeford et al. 2016) for weighted networks. The weighted clustering coefficient was calculated using the measure by (Onnela, Saramäki et al. 2005) and a threshold of ϕ_T_ = 0.6 was set for the small-world propensity to classify networks as small-world, according to previous described (Muldoon, Bridgeford et al. 2016). The SWP was normalized so that values closer to 1 indicates that the network is more small-world.

## 3 Results

### 3.1 AAV2/1 Gi-DREADD expression found exclusively in CaMKIIa positive neurons with maturing excitatory and inhibitory synapses

Immunolabeling with antibodies specific for CaMKIIa expressed on excitatory neurons, and for mCherry expressed by DREADDs confirmed that both proteins colocalize in the network as shown in **(Figure 2A)**. In addition, there was no colocalization with mCherry and GAD65/67, a marker for inhibitory neurons **(Figure 2B)**. Furthermore, at 14 DIV the networks expressed NMDA receptors **(Figure 3A)** and GABA **(Figure 3B)** indicating the capacity for excitatory and inhibitory signaling at this stage.

**Figure 2.**
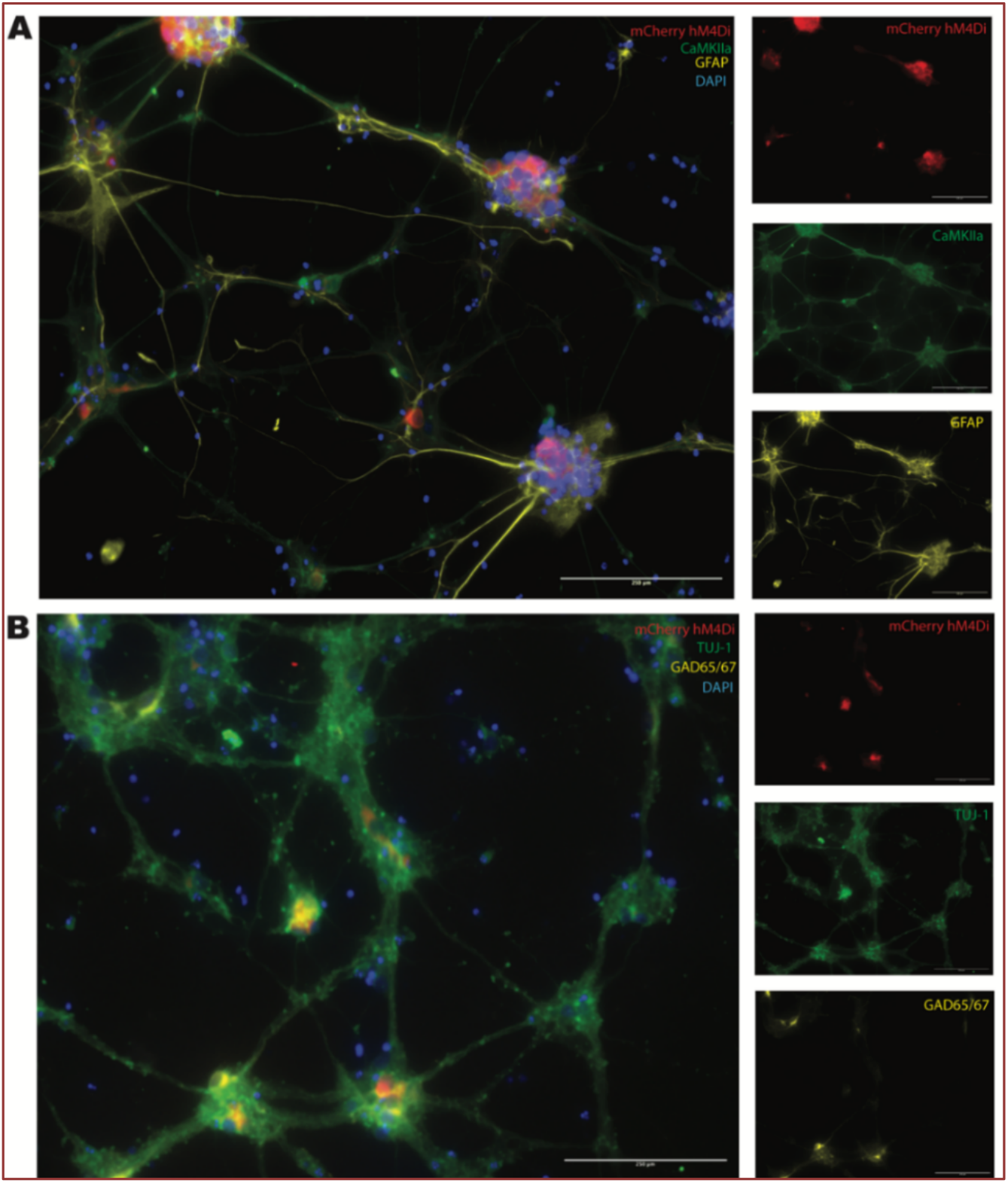
AAV2/1 hM4Di designer receptor exclusively activated by designer drugs (DREADDs) expressed in CaMKIIa positive neurons in vitro. **(A)** mCherry DREADDs expression was confirmed in CaMKIIa positive neurons **(B)** GAD65/67 expressing neurons (inhibitory neurons) did not express mCherry DREADDs. Scale bar = 250μm.

**Figure 3.**
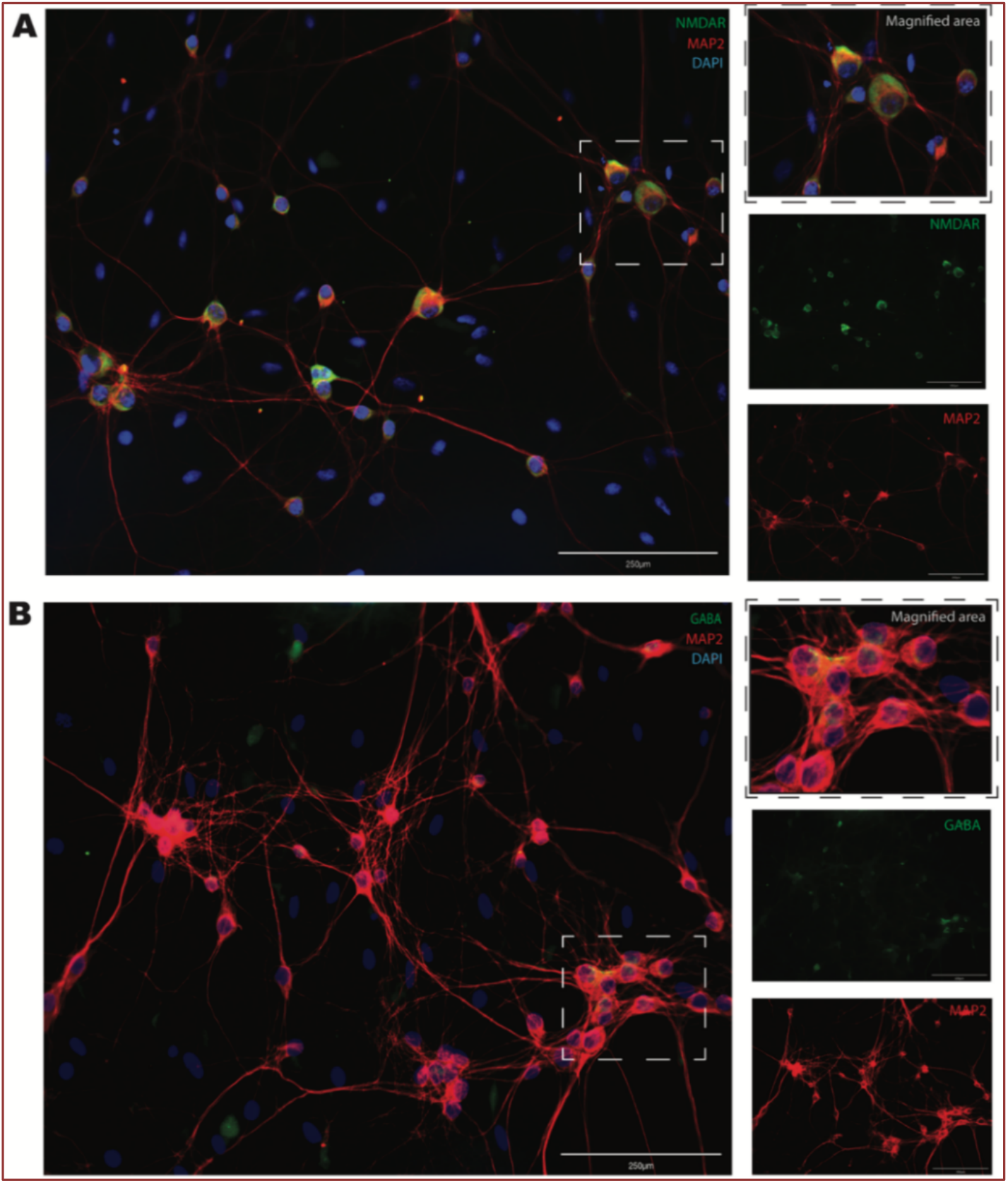
Neural networks positively stained for both NMDA receptors **(A)** and GABA **(B)** together with MAP2 neuronal cytoskeletal marker at 14 DIV. Scale bar = 250μm.

### 3.2 Network modularity develops over time and can be altered with inhibition of excitatory transmission

Spontaneous network activity was recorded between 14 DIV and up until 40 DIV. Graphs were created from the processed data to identify the different modules in the network and the nodes belonging to each module. These graphs are presented in **Figure 4** and show the modules for one network at 6 recording timepoints. At 14 DIV, the neural networks were still relatively immature and developing both structurally and functionally. Our results showed no distinct modules, and only a few active nodes were detected **(Figure 4A)**. At 21 and 26 DIV, almost all active nodes belonged to one of three modules detected **(Figure 4B, C; red, blue and yellow)**. The graph shown at 28 DIV **(Figure 4D)** depicts the modular organization in the network 24 hours after selective inhibition of excitatory transmission. The original 3 main modules from 26 DIV appeared to be fragmented into smaller clusters, with large areas where no activity was detected. Interestingly though, we found that by 32 DIV the network had begun reconfiguring itself, with more nodes organized into modules. By 40 DIV, the network had reconfigured itself back into 3 main modules, although with a different spatial positioning **(Figure 4E, F)**. Network organization at 40 DIV also showed that detected nodes within modules had more spatial distance compared to being positioned closer together at 21 DIV and 26 DIV.

**Figure 4.**
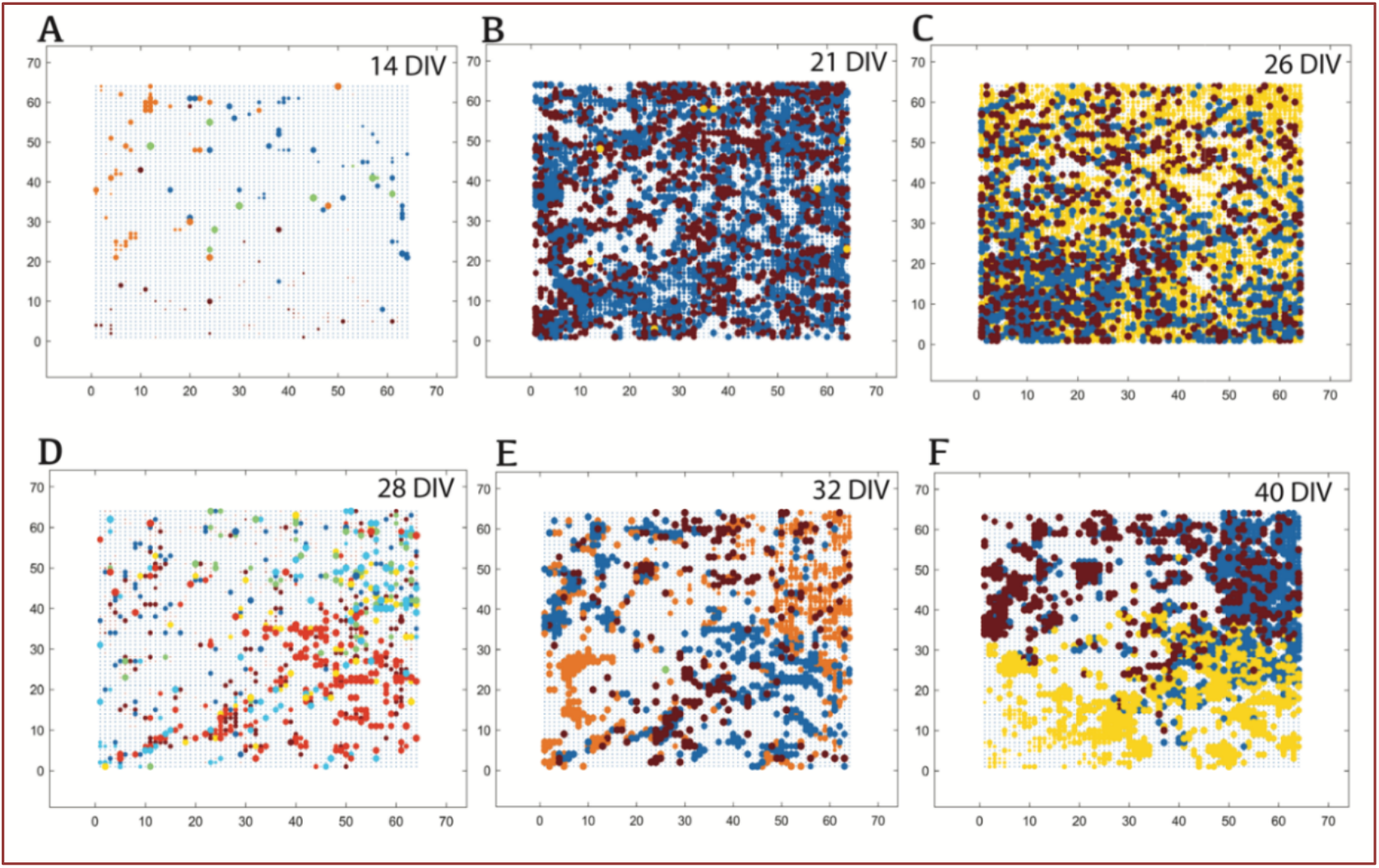
Inhibition results in functional de-clustering in modules with subsequent reorganization following recovery. Depicted here is a sample neural network across development. Modularity in the network at 14 DIV **(A)**, at 21 DIV **(B)** and at 26 DIV **(C)**. Network modular organization after perturbations at 28 DIV **(D)**, at 32 DIV **(E)** and at 40 DIV **(F)**. Each dot is one node and represents one active electrode at the time of recording. The colors indicate the module that the detected nodes belong to, and node sizes are scaled according to their participation coefficient.

### 3.3 Neural networks exhibited alterations in functional organization during and following selective perturbation

Further analyses were conducted to determine the extent to which both structural and functional organization had changed within the inhibited networks. **Figure 5** depicts the results of these analyses for 3 experimental networks, from here onwards identified as Net1, Net2 and Net3 (see figure legend). We observed inconsistencies in the number of modules before and after selective inhibition of the networks, making it difficult to conclude exactly the structural changes taking place. For instance, in **Figure 5A**, networks had self-organized into 3 modules at 21 DIV before inhibition. During the period of inhibition between 25 and 27 DIV, there was no change in the number of modules in Net2, an increase in module number from 3 to 6 modules at 27 DIV for Net1, and a decrease from 3 to 1 module at 27 DIV for Net3. During the network recovery period after inhibition, no modules were detected for Net2. The number of modules increased in Net1 and Net3, then returned to baseline number of 3 modules by 40 DIV.

**Figure 5.**
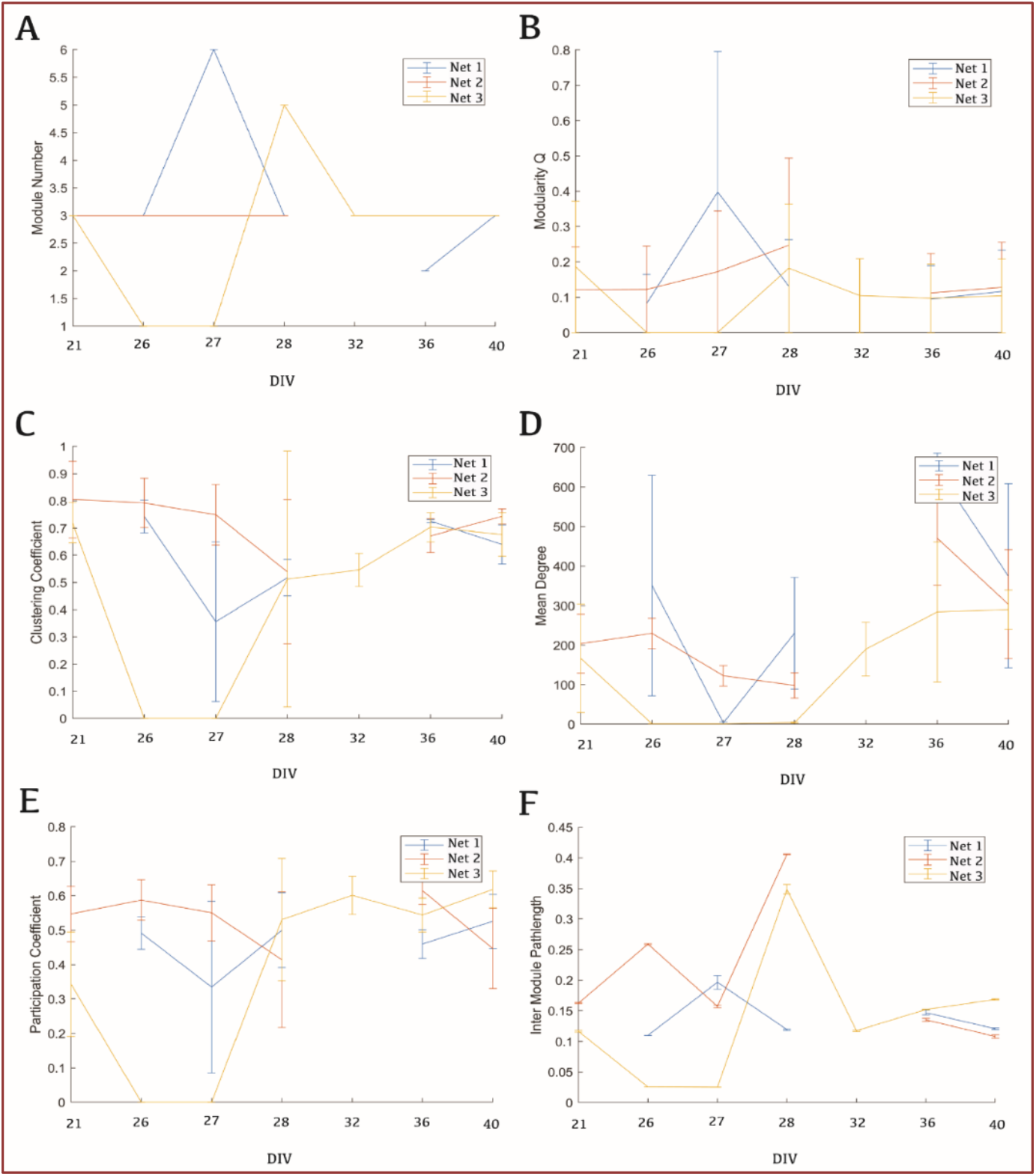
Neural networks exhibited transiently increased path lengths and decreased clustering due perturbation, with subsequent reconfiguration. Functional organization is described in terms of Module number **(A)**, Modularity Q **(B)**, Clustering coefficient **(C)**, Mean degree **(D)**, Participation coefficient **(E)**, and Inter module path length **(F)** with standard error bars.

For further analyses of network subdivision, analyses of the modularity Q were conducted, and the results highlighted that network organization reflected a community structure. Prior to perturbation, modularity Q = / > 0 but less than <0.2 **(Figure 5B)** indicating a community structure. Modularity Q appeared to increase (Q > 0.2) during perturbation for two networks (Net1 and Net3). One network decreased to Q = 0, indicating that there was low division during perturbation. Following perturbation and between 28 and 40 DIV, all networks appeared to stabilize with modularity Q values Q = 0.17 **(Figure 5B)**, which indicate varying modularity prior, during and after perturbation. Additional findings demonstrated that during the period of inhibition between 25 and 27 DIV, there was a decrease across all networks in clustering coefficient **(Figure 5C)**, however, all networks restored high clustering with (values > 0.6) after perturbation suggesting some recovery of network functional organization. Similar results were observed for the participation coefficient **(Figure 5E)**. Furthermore, the average number of connections that each node had with other nodes in the network had decreased from above >150 connections to <110 connections for Net2 during perturbation, while the other 2 networks had no detected connections by 27 DIV. However, during the recovery period after inhibition, the networks gradually restored the connections such that all networks had mean degrees of approximately > 300 connections by 40 DIV **(Figure 5D)**. Our examination of intermodular path length showed that the number of edges connecting modules fluctuated greatly between recordings during the period of inhibition between 25 and 27 DIV, as well as immediately after inhibition at 28 DIV **(Figure 5F)**. Nevertheless, we observed that networks restored to pre-perturbation values between 32 and 40 DIV at < 0.2 intermodular path length.

Finally, we found that the neural networks exhibited relatively large SWP values at 21 DIV (ϕ_T_ > 0.5) indicating that the neural networks displayed small-world properties before perturbation **(Figure 6)**. During perturbation, two networks maintained high SWP values, while one network decreased to ϕ_T_= 0.3. There were fluctuations in values during recovery, but by 40 DIV, all networks restored to SWP values ϕ_T_ > 0.5 although none restored to pre-inhibition values.

**Figure 6.**
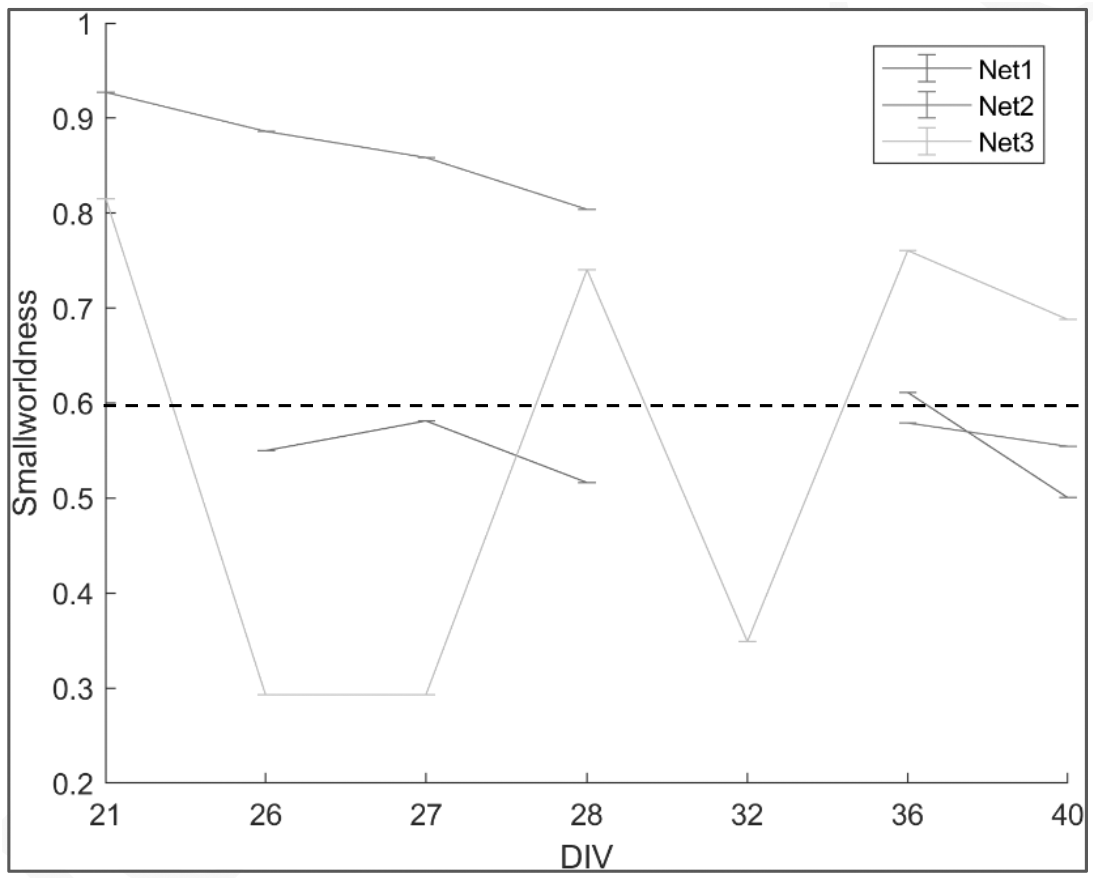
Neural networks exhibited transient decreases in SWP during perturbation, with subsequent recovery. The neural networks were analyzed as a weighted matrix with the dashed ling denoting the ϕ_T_ = 0.6 threshold.

## 4 Discussion

Complex biological systems like the brain or *in vitro* neural networks can adjust their structural and functional organization in response to changes in sensory inputs or experiences. In healthy conditions, network topological architecture evolves in accordance with the principles of high efficiency of information transfer and processing. Networks achieve this by having segregated modules for specialized processing, and short pathways to connect the different areas of the network (van den Heuvel and Sporns 2013). It is also imperative that the relevant topologies are resilient i.e., they need to be adaptable to perturbations. The findings presented in this study demonstrate that perturbation of network activity via selective inhibition of excitatory synaptic transmission resulted into transient de-clustering of modular structures that corresponded to reduced clustering and decreased small-worldness. Other structural changes included a transient increase in path lengths during perturbation, with the network subsequently reorganizing to a state of increased clustering and short path lengths following recovery.

By structurally and functionally reorganizing information processing areas in response to impaired synaptic transmission, complex biological networks can adapt to altered inputs (Rezaul Karim, Proulx et al. 2021) and retain function. De-clustering refers to the process by which specialized brain regions or modules lose their distinct functional boundaries. It involves a reduction in the functional segregation or modular organization of the brain, leading to increased interactions and information flow between previously distinct brain regions (Park and Friston 2013, Joanna Su Xian, Kwun Kei et al. 2019). Several studies have shown that decreased clustering can occur in various contexts including during brain development and aging (Micheloyannis, Vourkas et al. 2009, Joanna Su Xian, Kwun Kei et al. 2019). De-clustering also represents a dynamic process of functional reorganization that allows for more flexible and efficient information processing during perturbation, learning and sensory input by enabling new information to be integrated into existing ensembles (Katori, Sakamoto et al. 2011, Pinotsis, Brincat et al. 2017). Our previous investigation showed that selective inhibition results in an increase in network wide bursts and synchrony (Weir, Christiansen et al. 2023), which is highly relevant here because it suggests that global activation increases as inhibited networks lose their structural boundaries. This also suggests that the transient loss of functional boundaries between processing areas in the network may be a compensatory response to restore synaptic drive with and across the perturbed areas of the network. As synaptic inputs to different parts of the network decrease due to inhibition, the remaining areas may have increased their synaptic capabilities, for example by either scaling up neurotransmitter release, increasing receptors on neurites or lowering the threshold for excitatory post-synaptic current (Bridi, de Pasquale et al. 2018). As these synaptic modifications tend to occur on a slow time scale (Bridi, de Pasquale et al. 2018, Hobbiss, Ramiro-Cortés et al. 2018), it is entirely possible that they happen concomitantly with the increase in network synchrony since synchrony increases also occur gradually over several days (Weir, Christiansen et al. 2023).

Furthermore, the decrease in clustering coefficient and participation coefficient – two basic measurements of network communication and information flow – correspond with the reduction in the small-worldness in the network during and after inhibition. There are various factors that can influence small-worldness in neural networks, one of which is the lesioning of specific hubs. Although our perturbation did not target any particular area of the network, the observed effects closely align with other studies that have demonstrated the distinct impact of hub lesioning on the small-world structure of the remaining network (Sporns, Honey et al. 2007). Specifically, lesioning provincial hubs disturbs functional integration of the module to which they belong, which results in less segregation in the remaining network (Sporns, Honey et al. 2007). Evidently, by silencing excitatory synaptic transmission across the network, we also impaired hub function, which resulted in overall less discriminate processing. Provincial hubs play a pivotal role in functional processing in their module, while other hubs i.e., connector hubs, act as central communication transmission ports that receive and transfer a substantial bulk of information to the rest of the network (Bettencourt, Stephens et al. 2007). Since these hubs also make transmission faster by connecting several areas of the network, loss of hub processing will inevitably lead to some functional re-organization of paths in the network.

The findings presented in **Figure 5D, and F** indicate that the applied perturbation resulted in a decrease in mean degree, which implies a reduction in the average number of connections between nodes in the network. Additionally, the inter-module path length increased, indicating that transmission of signals between different areas of the network required more steps or intermediaries. These changes in network connectivity and information transmission have implications for the network’s efficiency. According to the theory of brain efficiency and economic cost of signal propagation (Achard and Bullmore 2007, Bullmore and Sporns 2012), a more efficient network is characterized by high degree of connectivity with direct information flow between nodes. A shorter path length between communicating areas of the network is also associated with improved efficiency as it minimizes the number of transmission steps required.

As with all complex self-organizing systems, there is an innate desire to self-organize towards increased effectiveness. This drive was evident in our study, where we observed the remarkable ability of neural networks to reconfigure themselves following perturbations at 25 DIV, 26 DIV and 27 DIV. By 32 DIV, the networks had already begun re-configuring to restore their segregated processing and distinct modularity **(Figure 4)**. Furthermore, the networks also exhibited the restoration of high mean degree and low path lengths, which would contribute to overall more efficient information processing, transmission, and communication. Together, these findings highlight the dynamic flexibility and self-reglatory capabilities of neural networks.

## 5 Conclusions and future directions

In conclusion, the findings from this study shed light on the remarkable adaptability of neural networks in response to perturbation, both structurally and functionally. We showed that targeted inhibition of excitatory synaptic transmission resulted in a temporary disruption in modular organization, network clustering and short path lengths. The neural networks also demonstrated an innate capacity to adapt and reconfigure themselves in response to perturbation, with the ultimate goal of restoring their functional organization and optimized information processing. Understanding the self-regulatory capabilities of neural networks provides valuable insights into the mechanisms underlying neural plasticity, and recovery from disruptions. Therefore, our study highlights the significance of utilizing *in vitro* models as tools to explore these intricate structure-function relationships at a micro and mesoscale level in changing conditions in complex neural networks. Future analyses to validate and characterize other features of structural and functional organization such as, hub organization, degree distribution and specific measure of efficiency may help to strengthen the findings reported here. These further studies would also benefit greatly from expanding the sample size and data set to investigate network resilience at different timepoints in development i.e., early development and late maturity. Together, the results can enhance our understanding of adaptability in neural networks and may lead to the development of therapeutic strategies targeting neuroplasticity for function restoration in various neurological conditions.

## Data availability statement

The raw data that support the findings of this study will be made available by the authors upon request.

## Author Contributions

This study was conceived and designed by JSW, AS and IS. All data collection was done by JSW who also drafted the manuscript. OHR performed all data analysis. AS and IS helped with the manuscript revision process, and all authors have read and approved the final version of the manuscript.

## Funding

This project was funded by The Research Council of Norway (NFR IKT Pluss; Self-Organizing Computational Substrates (SOCRATES)) Grant number: 270961,

## Acknowledgement

The authors would like to thank Dr. Rajeevkumar Nair Raveendran at the Viral Vector Core Facility, Kavli Institute for Systems Neuroscience for designing and manufacturing the AAV vectors.

## Conflict of Interest

The authors declare that the research was conducted in the absence of any commercial or financial relationships that could be construed as potential conflict of interest.

## Notes

### Competing Interest Statement

The authors have declared no competing interest.

